# The switch from bacterial phosphorus mineralization to arbuscular mycorrhiza in root hairless wheat during crop development

**DOI:** 10.1101/2025.08.26.672279

**Authors:** Courtney Horn Herms, Ian Tsang, Frederik Bak, Klara Gunnarsen, Magdalena Hasenzagl, Amaru Miranda Djurhuus, Tom Thirkell, Eric Ober, Fiona Leigh, James Cockram, Mette Haubjerg Nicolaisen

## Abstract

Understanding how crops recruit bacterial and fungal partners for phosphorus (P) nutrition remains incomplete, particularly regarding temporal shifts between P-cycling bacteria and arbuscular mycorrhizal fungi (AMF). We investigated these dynamics using the wheat root hair mutant *short root hair* 1 (srh1), which is impaired in root hair elongation and may be more dependent on microbial P acquisition compared to the wildtype. We profiled the P-cycling bacterial microbiome and AMF colonization across development and at booting under P-depleted conditions. Before AMF establishment, the root hair mutant rhizosphere was enriched in bacterial organic P-mineralization genes. After AMF colonization increased in both genotypes, this enrichment disappeared, indicating a temporal shift from bacterial to fungal P acquisition. Growing the root hair mutant in P-depleted conditions to booting, when P uptake is highest in wheat, did not induce the mutant to recruit its bacterial P-cycling microbiome. Instead, AMF arbuscule formation was enriched. Collectively, our results reveal a temporal transition in wheat from bacterial to AMF-mediated P nutrition and highlight the compensatory role of the microbiome in plants lacking root hair elongation.

## Introduction

Agricultural productivity is often constrained by nutrient availability. Phosphorus (P) is an essential nutrient and vital for central biochemical processes in the plant (Rengel et al., 2022). Despite relatively large amounts of total P in most agricultural soils, bioavailability of P is limited, leading to restricted plant growth and yield (Menezes-Blackburn et al., 2018). Thus, P is one of the most heavily applied nutrients to agricultural fields. P fertilizer is, however, poorly absorbed by the plant (Simpson et al., 2011), leading to risks of environmental runoff and eutrophication (Ulén et al., 2007). Furthermore, raw phosphate rock used to make P fertilizer is a finite resource with high energy demands for production (Van Kauwenbergh, 2010), further impacting the sustainability of current P fertilization practices. The need to grow more food using reduced P inputs requires the development of new methods to improve crop P nutrition under reduced chemical fertilizer amendments.

The plant utilizes several methods to ensure P nutrition. Root hairs help the plant access greater volumes of soil and improve P uptake (Bates & Lynch, 1996; Gahoonia et al., 2001). In addition, the plant root microbiome plays an important role in plant P nutrition. The fungal fraction of the microbiome, notably the arbuscular mycorrhizal fungi (AMF), play a vital role in accessing soil P and providing mineral P to their host plant, including in globally important crops such as wheat (*Triticum aestivum* L.) (Thirkell et al., 2020). In low-P conditions, AMF colonization is enriched to improve crop P uptake (Breuillin et al., 2010). Similarly, in root hairless crops, AMF take over the role of root hairs in P uptake in accordance with the root economics spectrum (Kumar et al., 2019; Wang et al., 2025). AMF colonize plant roots at a defined point in root development; in field-grown wheat, this correlates to growth stage (GS) 12 (Marrassini et al., 2024) on Zadok’s scale of wheat development (Zadoks et al., 1974), where two leaves are fully unfolded.

The bacterial fraction of the plant root microbiome is initially recruited from the seed-borne microbial community (Simonin et al., 2022). From the seed and throughout germination and development, the plant develops its bacterial microbiome to assist in P cycling (Xiong et al., 2021; Nunes et al., 2022) and to alleviate P stress when P uptake is limited (Tang et al., 2022; Zhang et al., 2025). Organic P mineralization is represented by bacterial genes such as *bpp* (beta propeller phytase), *phnK* (phosphonate degradation*), phoD* (cytoplasmic phosphatase), and *phoX* (periplasmic phosphatase). Bacterial solubilization of inorganic P can occur through acidification via gluconic acid, the production of which requires the specific co-factor pyrroloquinoline-quinone (PQQ). However, bacteria receive little attention in the root economic space (Matthus et al., 2025), and especially in regards to P nutrition. Unraveling how plants integrate both rhizosphere bacteria and AMF to address their P needs may inform strategies to leverage plant-microbe interactions for more sustainable agriculture.

AMF colonize plant roots only when the root system is sufficiently developed (Werner & Kiers, 2015), whereas P-cycling bacteria are present from early seed germination (Nunes et al., 2022). However, little is known about the enrichment of P-cycling bacteria in the rhizosphere before and after AMF colonization. To address this gap in knowledge, we compared the P-cycling bacterial microbiome between the recently characterized wheat root hair mutant *short root hair 1* (*srh1*) and its wild-type background (Tsang et al., 2024). We hypothesized that the wheat root hair mutant enriches P-cycling bacteria to a greater extent than its wild-type background in early development, before the AMF symbiosis is prevalent, to compensate for the lack of root hairs. After the AMF symbiosis is established, we hypothesized that the root hair mutant and the wild-type will have similar bacterial P-cycling levels, due to the dependency on AMF for P nutrition. Furthermore, we hypothesized that we could stimulate the root hair mutant to retain its enriched P-cycling microbiome to later stages of its development by growing the plants in P-depleted conditions. To address these hypotheses, we harvested the root hair mutant and its wild-type background from the field at GS11 and GS13 to capture the response of the P-cycling bacterial community before and after AMF establishment. To evaluate the AMF and bacterial fractions of the P-cycling microbiome at later developmental stages, we harvested a set of plants grown in pots to booting (GS45-47), at which time P uptake is highest in wheat. In this pot-based setup, we could also manipulate the P status of the soil. Long-read 16S ribosomal RNA (rRNA) amplicon sequencing complemented quantitative polymerase chain reaction (qPCR)-based functional analyses of the bacterial microbiome, while AMF were quantified through qPCR and traditional staining.

## Materials and Methods

### Plant material

The wheat root hair mutant, *srh1,* and its corresponding wild-type line have been characterized in full (Tsang et al., 2024). Briefly, the mutant was identified in a forward screen of an ethyl methanesulfonate-mutated population created in the spring wheat cultivar Cadenza (Krasileva et al., 2017). In the root hair mutant, root hairs initiate but fail to elongate and were measured to be approximately 0.15 mm long compared to the wild-type length of approximately 1.90 mm (Tsang et al., 2024).

### Field experiment set-up and sampling

Wild-type and mutant seeds were sown in spring 2024 in a complete randomized block design in Hinxton, Cambridgeshire, United Kingdom (52°05’56.1”N, 0°10’41.8”E). The soil pH, Olsen extractable P, and ammonium nitrate extractable potassium (K) and magnesium (Mg) were measured on unplanted soil by NRM Cawood Scientific (Bracknell, UK) (Table S1). Approximately 120-180 seeds were sown per plot, where each plot measured 1x1 m with 6 rows of plants per plot. Plants were excavated at GS11 and GS13 (Zadoks et al., 1974) (two and four weeks post-sowing, respectively). One individual plant was sampled from each replicate plot (n = 10) using a sterilized shovel, with care to harvest the entire root system.

Plants were gently shaken to remove the loosely attached bulk soil. The shoot length and the length of the longest root of the root system were measured. The rhizosphere sample (soil influenced by the root) was taken following the procedures described by Guan et al. (2024). Briefly, the roots were placed in 25 mL sterile Milli-Q water and shaken for 30 s; the resulting soil was collected as rhizosphere soil. At each sampling time point, bulk soil (n = 5) was also collected from the bare soil between the plots. The rhizosphere and bulk soil samples were stored at -80 °C until freeze-drying under vacuum at -70 °C. Thereafter, the dried soil samples were stored at 4 °C until subsequent DNA extraction. Shoots were dried at 37 °C, and the shoot dry weight was recorded.

Roots were stored in 70% ethanol at 4 °C prior to root scanning. Root scans were taken using an Epson Dual Lens System Epson Perfection V800 Photo at 600 DPI in greyscale. RhizoVision Explorer version 2.0.3 (Seethepalli et al., 2021) was used to analyze the root scans to calculate total root length, average diameter, perimeter, volume, surface area, and branching frequency. Scans were taken in “Broken roots” mode and at an image thresholding level of 175. Edge smoothing was enabled with a threshold of 3, and root pruning was enabled at a threshold of 10. Non-root objects below 2 mm^2^ were removed. After root scanning, the roots were dried at 37 °C, and the dry weight was subsequently recorded.

### Pot experiment set-up and sampling

High- and low-P field soil was collected from the Sawyers Experimental Field site at Rothamsted Research (Harpenden, UK; 51° 48′ 59.39″ N, 0° 22′ 33.28″ W). Soils from two historically low-P plots (last measured Olsen P = 7.1 mg/kg and 5.7 mg/kg in February 2022; A. Riche, personal communication, May 1, 2024) and two historically high-P plots (last measured Olsen P = 19.3 mg/kg and 18.6 mg/kg in February 2022; A. Riche, personal communication, May 1, 2024) were collected and mixed to create the low- and high-P potting soil, respectively. The soil was sieved through 4 mm mesh and mixed with washed sand (Horticultural Silver Sand, Melcourt, Tetbury, UK) in a 3:2 ratio (w/w) to improve drainage and root aeration. Samples of the soil-sand mixture were analyzed for pH, Olsen extractable P, and ammonium nitrate extractable potassium (K) and magnesium (Mg) by NRM Cawood Scientific (Bracknell, UK) (Table S1).

Two-liter pots (12 cm H x 12 cm W x 20 cm L) were sterilized (Hortisept Pro, Conka, Nelson, UK), dried, and filled with the soil-sand mixture. Wild-type and mutant seeds (n = 8) were pre-germinated on damp filter paper in the dark for three days before planting one seed per pot. Plants were grown in a climate chamber (Conviron CMP6010) under 16-hour days at 20 °C, 60% humidity, and light level 3, followed by 8-hour nights at 18 °C and 60% humidity. Throughout growth, pots were rotated twice a week and watered as needed approximately three times a week. Plants were harvested at booting (GS45-47; Zadoks et al., 1974) after five weeks of growth. The leaf sheath of the plants was swollen and just started to open to expose the ear.

Plants were gently removed from the pots, and the root system was divided in half along the longitudinal axis. Approximately half the root system was used for rhizosphere sampling as described in Guan et al. (2024). Briefly, the roots were placed in 25 mL sterile Milli-Q water and shaken for 30 s; the resulting soil was collected as rhizosphere soil. Rhizosphere soil samples were stored at -80 °C until freeze-drying under vacuum at -70 °C. Thereafter, the dried rhizosphere soil samples were weighed as a proxy for rhizosheath size and stored at 4 °C until DNA extraction. Shoots were dried at 37 °C and subsequently the dry weight was recorded. Roots were stored in 50% ethanol at 4 °C until AMF quantification.

Dried shoots were homogenized and ground using a zirconium ball mill, and the P concentration was determined using the dry ashing method of Saunders & Williams (1955) followed by the molybdate-blue method (Murphy & Riley, 1962) using flow injection analysis (FIAstar 5000 analyzer, Foss, Hillerod, Denmark). Briefly, 100 mg samples were placed in acid-washed crucibles and ashed in a muffle furnace at 550 °C for one hour. The ash was mixed with 50 mL 0.5 M sulfuric acid and shaken overnight at room temperature. Extracts were filtered through ashless filters (Whatman No. 42), and samples were diluted 10X before analysis. Two blanks and one replicate of the standard reference material 1515 (NIST 1991) (Sigma Aldrich, Denmark) were included in the analysis.

### AMF quantification

Due to the small and fragile root system of the young plants at GS11 and GS 12-13, relative qPCR was used to quantify AMF colonization (Bodenhausen et al., 2021). DNA was extracted from the dried roots using a FastPrep-24 5G beadbeating system (MP Biomedicals, Irvine, CA, USA) at 6.0 m/s for 40 s and the FastDNA SPIN Kit (MP Biomedicals, Irvine, CA, USA) following the manufacturer’s instructions. DNA quality and quantity were recorded using a NanoDrop ND-1000 spectrophotometer (Thermo Fisher Scientific, Carlsbad, CA, USA) and a Qubit 2.0 Fluorometer (Thermo Fisher Scientific, Carlsbad, CA, USA), respectively. DNA was diluted ten-fold to a concentration between 1 and 10 ng/μL. AMF colonization was quantified using the 18S rRNA gene fragment, amplified with the AMF-specific primers AMG1F (Hewins et al., 2015) and AM1 (Helgason et al., 1998) (Table S2) on an AriaMx Real-Time PCR system (Agilent Technologies, Santa Clara, CA, USA) using the PCR conditions described in Bodenhausen et al. (2021). The AMF 18S rRNA gene signal was normalized using a wheat gene encoding a heterogenous nuclear ribonucleoprotein Q, *hn-RNP-Q*, which was amplified with previously described primers (Table S2) and PCR conditions (Qi et al., 2012). For both primer sets, each 20 µL qPCR reaction contained 10 µL 2X Brilliant III Ultra-Fast SYBR Green qPCR Master Mix with Low ROX (Agilent Technologies, Santa Clara, CA, USA), 0.4 µM each of forward and reverse primer, 1 mg/mL bovine serum albumin (New England Bioabs, MA, USA), and 2 µL of template DNA. qPCR products were visualized on a 1% agarose gel using GelRed (Sigma Aldrich, Denmark) to confirm amplification of the target gene. Negative controls were run in triplicate for each primer pair. Raw fluorescence data were imported to LinRegPCR (version 2021.2) (Ruijter et al., 2009) to determine cycle number to threshold (*Ct*) and efficiency (*E*) using the default baseline threshold from LinRegPCR. The AMF 18S rRNA gene signal was normalized to plant DNA using the formula 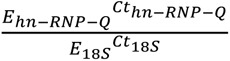 as described by Bodenhausen et al. (2021).

At booting (GS45-47), AMF colonization was quantified via staining (Phillips & Hayman, 1970). From each plant, approximately 0.2 g of root taken from the middle of the root system was stained with Trypan Blue. Root samples were incubated in 2 mL 10% KOH for 25 min at 90°C. Subsequently, the KOH was removed, and the roots were washed twice with distilled water. After washing, roots were incubated in 2 mL 0.3 M HCl for 30 min at room temperature. After removing the HCl, roots were incubated in 2 mL 0.1% Trypan Blue stain for eight minutes at 90°C. Roots were then washed twice with acidic glycerol (1:1 glycerol:0.3 M HCl v/v) and incubated with the last wash overnight at room temperature in the dark. At least ten stained root sections measuring approximately 1 cm per sample were mounted on microscope slides. Scoring for AMF presence and arbuscule formation was performed on 100 fields of view per slide using a Leica DMLS microscope (Leica Microsystems, Wetzlar, Germany) under 200X magnification (Brundrett et al., 1996).

### Bacterial P-cycling gene quantification and 16S rRNA gene amplicon sequencing

DNA from up to 500 mg freeze-dried rhizosphere samples was extracted using a FastPrep-24 5G beadbeating system (MP Biomedicals, Irvine, CA, USA) at 6.0 m/s for 40 s and the FastDNA SPIN Kit for soil (MP Biomedicals, Irvine, CA, USA) following the manufacturer’s instructions. DNA quality and quantity were tested using a NanoDrop ND-1000 spectrophotometer (Thermo Fisher Scientific, Carlsbad, CA, USA) and a Qubit 2.0 Fluorometer (Thermo Fisher Scientific, Carlsbad, CA, USA), respectively. DNA was diluted ten-fold to a quantity between 1 and 10 ng/μL.

Quantification of the 16S rRNA gene and bacterial P-cycling genes (*bpp*, *phnK*, *phoD*, *phoX*, and *pqqC*) in the rhizosphere samples was performed via qPCR using the AriaMx Real-Time PCR system (Agilent Technologies, Santa Clara, CA, USA) following the qPCR conditions described in Hao et al. (2020) and primers in Table S2. qPCR products were visualized on a 1% agarose gel using GelRed (Sigma Aldrich, Denmark) to confirm amplification of the target gene. Standard curves and negative controls were run in triplicate on each plate. Abundance of the P-cycling genes was normalized to 16S rRNA copies.

The bacterial community in the rhizosphere was analyzed by sequencing 1050 bp of the 16S rRNA gene, amplified using primers 341F (5’-CCTACGGGNGGCWGCAG -3’) and 1391R (5’-GACGGGCGGTGWGTRCA -3’) (Thijs et al., 2017) with the Platinum SuperFi II polymerase (Thermo Fischer Scientific, Waltham, USA). The ZymoBIOMICS Microbial Community DNA Standard (Zymo Research, Irvine, CA, USA) was included as a sequencing control. The PCR cycling conditions were as follows: an initial denaturation step of 98 °C for 30 s, followed by 30 cycles of denaturation at 98 °C for 10 s, annealing at 64 °C for 30 s, and elongation at 72 °C for 45 s, and a final elongation at 72 °C for 5 min. PCR products were visualized via electrophoresis on a 1% agarose gel before purification using AMPure XP beads (Beckman Coulter Inc., Brea, CA, USA) at a 0.6 ratio. The sequencing library was prepared using a native barcoding kit V14 (SQK-NBD114.96) and sequenced using a PromethION R10.4.1 flow cell on the PromethION P2 Solo platform (Oxford Nanopore, UK).

Adapter trimming and basecalling were performed using Dorado version 0.7.0 using the super accurate setting (github.com/nanoporetech/dorado). Seqtk (github.com/lh3/seqtk) was used to discard reads below 1000 bp and above 1500 bp. Amplicon sequences were extracted from between the primer sequences using the script strip_degen_primer_deep (github.com/padbr/asat). NanoFilt version 2.8.0 (De Coster et al., 2018) was used to discard amplicon sequences with an average quality score below 25. Cutadapt version 3.5 (M. Martin, 2011) was used to truncate longer reads at 1010 bp. The remainder of the analysis was performed in R version 4.3.1 (R Core Team, 2023). The reads were dereplicated to form amplicon sequence variants (ASVs) and assigned taxonomy using dada2 version 1.22.0 (Callahan et al., 2016) with the SILVA database version 138.2 (Quast et al., 2013). The ASV table was transformed to a phyloseq object for subsequent analysis using phyloseq version 1.44.0 (McMurdie & Holmes, 2013). Non-bacterial ASVs and those assigned to chloroplasts and mitochondria were discarded before the ASVs were agglomerated at the genus level while retaining non-classified reads. The ZymoBIOMICS Microbial Community DNA Standard (Zymo Research, Irvine, CA, USA) was used to assess sequencing contamination. Ampvis2 version 2.8.1 (Andersen et al., 2018) was used to visualize bacterial community composition. A dissimilarity matrix was calculated with rarefaction to 6639 reads using avgdist from the vegan package version 2.6-4 with default parameters (vegandevs.github.io/vegan) and was used to generate the principal coordinate analysis based on Bray-Curtis dissimilarities and to run a PERMANOVA using adonis2 from vegan. Pairwise comparisons were made with the package pairwiseAdonis version 0.4.1 (github.com/pmartinezarbizu/pairwiseAdonis). The Shannon diversity index (alpha diversity) was calculated 100 times, as proposed by Schloss (2024). Briefly, the dataset was rarefied to 6639 reads, and the Shannon diversity index was subsequently calculated as the mean Shannon diversity index over 100 replicates. Differentially abundant ASVs between the mutant and wild-type were generated using corncob version 0.4.1 (B. D. Martin et al., 2020) with a false discovery rate of 0.05.

Raw reads are deposited in NCBI under BioProject PRJNA1308004.

### Statistical analysis

Data analysis was conducted in R version 4.3.1 (R Core Team, 2023), and all plots were created using ggplot2 version 3.5.1 (Wickham, 2016). Significance was defined as p < 0.05. When comparing means, the Shapiro-Wilk test was used to check for normality. The Student’s t-test was used to compare means for parametric groups, while the Wilcoxon test was used for non-parametric groups. All qPCR data were log-transformed before analysis, and due to heteroscedasticity and different data structures across growth stages, separate models were made for each gene in each growth stage. All analyses of variance (ANOVAs) were performed using the car package version 3.0-5 (Fox & Weisberg, 2019). All post-hoc pairwise comparisons were performed using the emmeans package version 1.10.4 (rvlenth.github.io/emmeans) with Tukey’s Honestly Significant Difference (HSD) test.

## Results

### Loss of root hair elongation impacts wheat development across growth stages

We initially evaluated the effect of the *srh1* mutation, which arrests root hair elongation (Tsang et al., 2024), on overall plant performance in real soils across three growth stages: first leaf fully unfolded (GS11), three leaves unfolded (GS13), and booting (GS45-47). At the two youngest growth stages, aboveground growth was no different between the two genotypes; the root hair mutant had longer shoots at GS11 (Student’s t-test, p < 0.05), but this was not reflected in shoot biomass (Student’s t-test, p > 0.05, Figure 1A-B). On the other hand, root growth was stunted in the root hair mutant at GS11 but restored at GS13 (Figure S1). The lack of root hair elongation restricted rhizosheath development across the growth stages (Student’s t-test, p < 0.05), although this was obscured at GS13 due to wet soils at harvest and enlarged rhizosheath size (Figure 1C).

**Figure 1.**
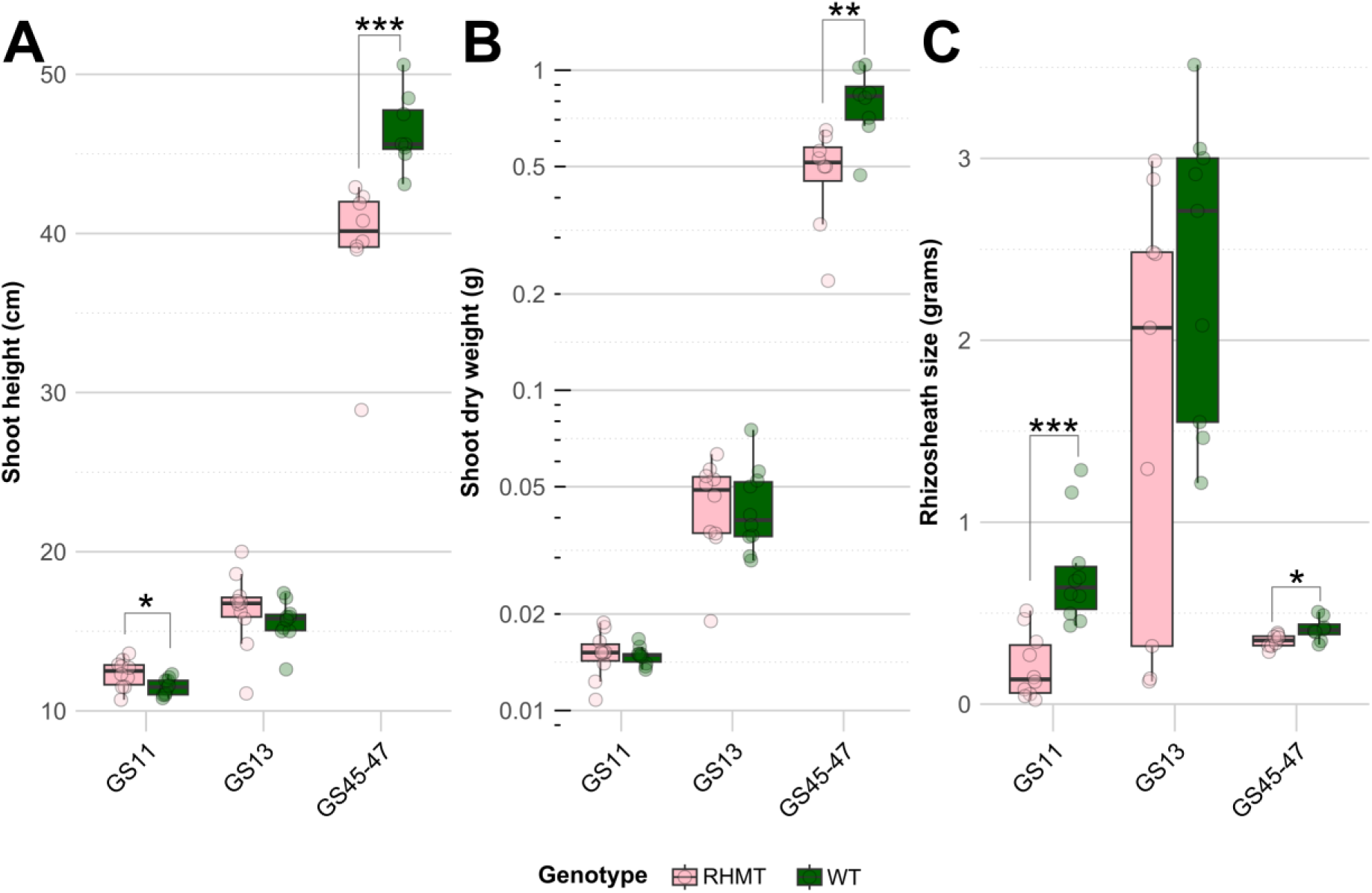
Plant performance of the root hair mutant (RHMT) and wild-type (WT), measured as shoot height (A), shoot dry weight (B), and rhizosheath size (C) across growth stages. Genotypes were compared within each growth stage using t-tests (parametric data) or Wilcoxon tests (non-parametric data). *p < 0.05, **p < 0.01, ***p < 0.001.

At booting (GS45-47), the phenotypic difference between the genotypes was apparent. The root hair mutant was stunted in aboveground growth compared to the wild-type (Student’s t-test or Wilcoxon test, p < 0.05, Figure 1A-B). However, shoot P concentration at this growth stage was not different between the root hair mutant (2.84 mg/g ± 0.374 mg/g) and the wild-type (2.47 mg/g ± 0.246 mg/g) (Wilcoxon test, p > 0.05).

### Initial higher colonization of bacteria harboring P mineralization genes in the wheat root hair mutant

The ability of the root hair mutant to enrich its bacterial microbiome for P-cycling was assessed alongside the development of the AMF symbiosis across the three growth stages.

At GS11, the relative abundance of all bacterial P-cycling genes, except for *phoX*, was affected by sample source (root hair mutant, wild-type, or bulk soil) at GS11 (one-way ANOVA, p < 0.05, Table S3). Bacterial genes involved in organic P mineralization (*bpp*, *phnK*, and *phoD*) were enriched in the rhizosphere of the root hair mutant compared to the wild-type (Tukey’s HSD, p < 0.05, Table 1). No differences in the relative abundance of *phoX* (periplasmic phosphatase) and *pqqC* (pyrroloquinoline-quinone synthase) were found between the root hair mutant and wild-type (Tukey’s HSD, p > 0.05). There was low AMF signal in both root hair mutant and wild-type plants, and the log2 ratio of AMF DNA to plant DNA was below zero. There was no enrichment of AMF signal in either genotype (Figure 2, two-way ANOVA, p > 0.05, Table S4).

**Figure 2.**
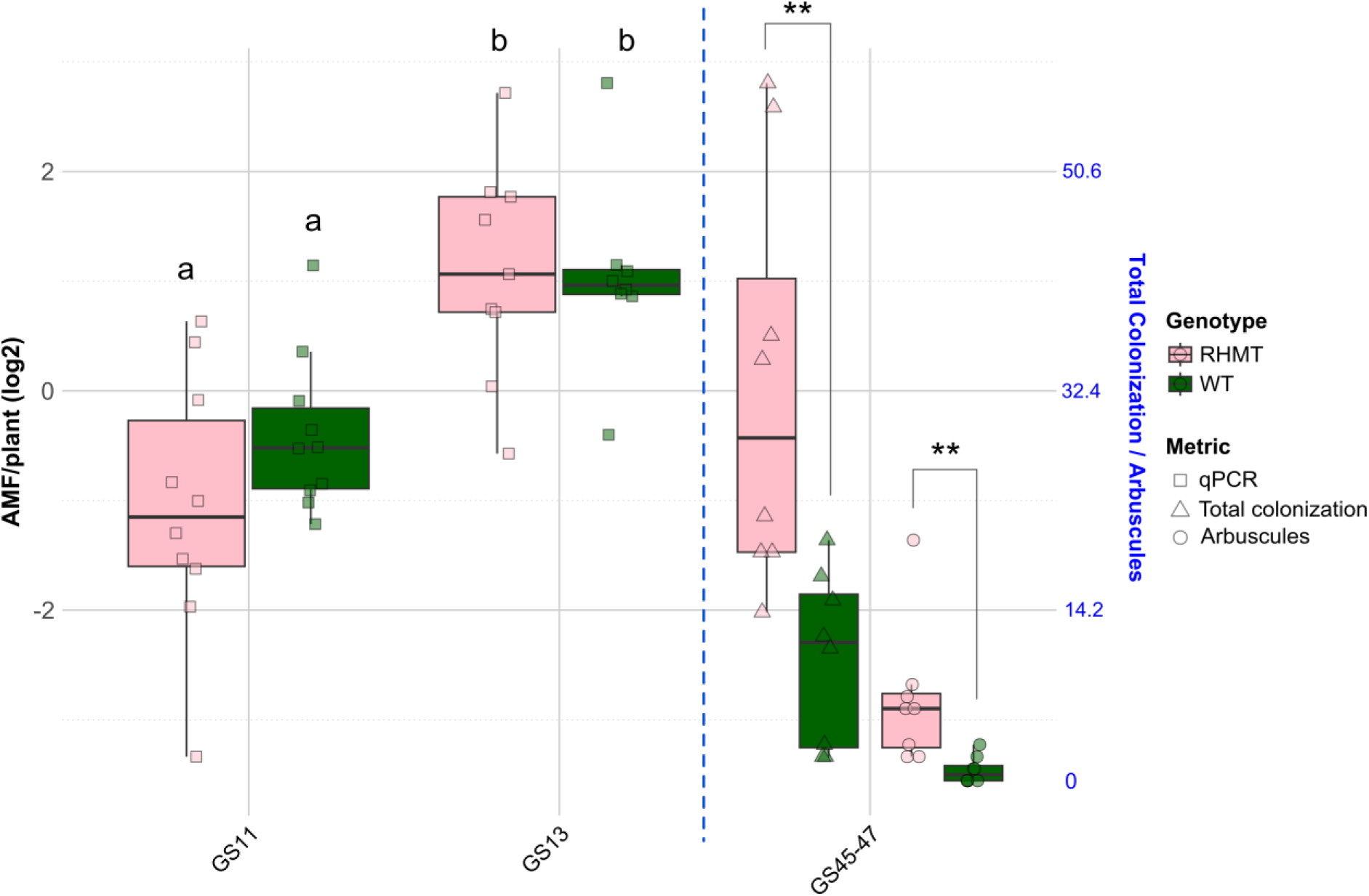
AMF signal in the root hair mutant (RHMT) and wild-type (WT) quantified via qPCR or via microscopy (percent total colonization and percent arbuscule formation) across the growth stages. The boxplots display the median, the two hinges (25th and 75th percentile), and the upper and lower whiskers (extending to the largest and smallest values no further than 1.5 times the interquartile range). True data points are overlaid on the boxplots as filled circles. qPCR means were compared using one-way ANOVA with Tukey’s HSD post-hoc test; means that share the same letter are not statistically different (p < 0.05). Microscopy means (percent total colonization and arbuscule formation) were compared with Wilcoxon test. **p < 0.01.

**Table 1.**
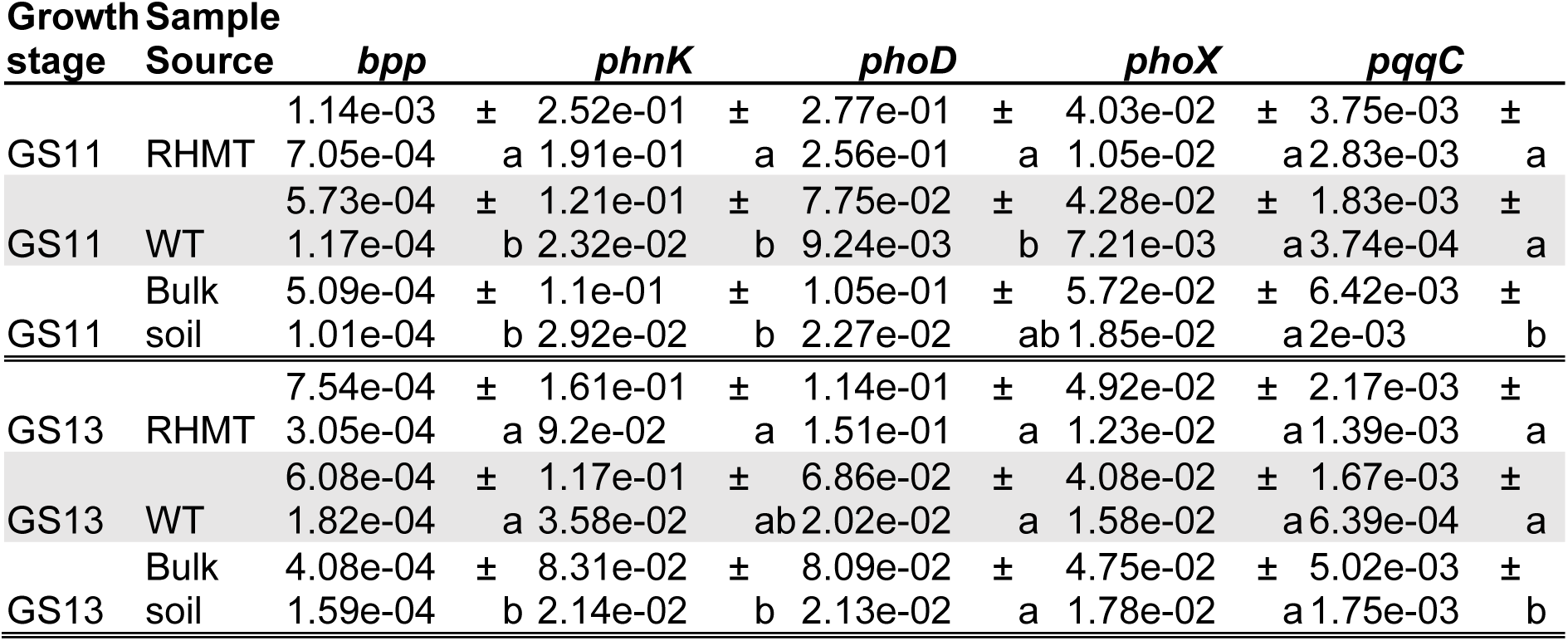
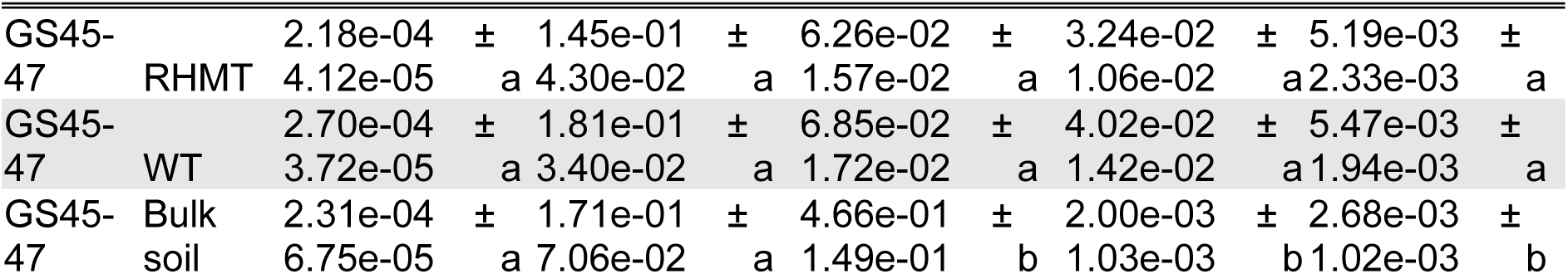
Relative abundance of bacterial P-cycling genes in root hair mutant (RHMT) and wild-type (WT) rhizosphere. Relative abundance was calculated by normalizing against 16S rRNA gene copies, and the mean is presented ± the standard deviation. Log-transformed gene relative abundance was compared among genotypes within each growth stage using one-way ANOVA with Tukey’s HSD post-hoc test. Means that share the same letter are not statistically different (p < 0.05).

At GS13, the relative abundance of *bpp*, *phnK,* and *phoD* in the rhizosphere of the root hair mutant decreased to the level of the wild-type. While sample source exhibited an effect on gene relative abundance (one-way ANOVA, p < 0.05, Table S3), with the exception of *phoD* and *phoX*, this was driven by the difference in gene abundance between the plant rhizospheres and bulk soil (Table 1). The decrease in P mineralization gene relative abundance coincided with an increase in AMF signal across the plants at this growth stage. The ratio of AMF DNA to plant DNA increased and was higher than at GS11, indicating the establishment of the AMF symbiosis (Figure 2; two-way ANOVA, p < 0.05, Table S4), but there was no effect of genotype on AMF signal (two-way ANOVA, p > 0.05, Table S4).

At the latest growth stage, GS45-47, there was no enrichment of bacterial P-cycling genes in the root hair mutant compared to the wildtype, similar to GS13 (Table 1, Tukey’s HSD, p > 0.05). Only the bulk soil was different from the plant rhizospheres for the genes *phoD*, *phoX*, and *pqqC* (Tukey’s HSD, p < 0.05). On the other hand, microscopy indicated increased total AMF colonization in the root hair mutant compared to the wild-type (Figure 2, Wilcoxon test, p < 0.05). Arbuscule formation reflected the result of total colonization, with the mutant exhibiting elevated arbuscule formation compared to the wild-type (Figure 2, Wilcoxon test, p < 0.05).

In summary, the results showed that at early stages of plant growth prior to AMF root colonization, a higher abundance of bacteria involved in P mineralization colonized the rhizosphere of the root hair mutant compared to the wild-type. This phenomenon was no longer observed upon establishment of AMF symbiosis, and bacterial P cycling genes were no longer enriched in the root hair mutant rhizosphere.

### Loss of root hair elongation had minor effects on bacterial taxonomic composition in the rhizosphere across development

To reveal the differences in rhizosphere bacterial community composition between the root hair mutant and the wild-type, long-read 16S rRNA amplicon sequencing was performed on the rhizosphere of the plants across the three growth stages. After filtering, 1,264,834 reads were retained. The median number of reads per sample was 14,570. 80% of the ASVs were identified at genus level.

Across the three growth stages, genera such as *Bacillus, Niallia, and Flavobacterium* were most abundant in both root hair mutant and wild-type plants (Supplemental Figure S2). There was no effect of root hair elongation on the alpha diversity of the bacterial microbiome in the rhizosphere across the three growth stages (Supplemental Figure S2, Wilcoxon test, p > 0.05).

No bacterial genera were differentially enriched in either genotype at GS11. On the other hand, at GS13, *Arthrobacter*, *Ensifer*, *Pseudomonas*, *Stenotrophomonas*, *Pedobacter*, and *Pantoea* were enriched in the root hair mutant, while *Ilumatobacter* and an unidentified genus of the Fibrobacteraceae family were enriched in wild-type plants (Figure 3A). At GS45-47, only an unclassified genus of the Mycoplasmataceae family was enriched in the root hair mutant (Figure 3B).

**Figure 3.**
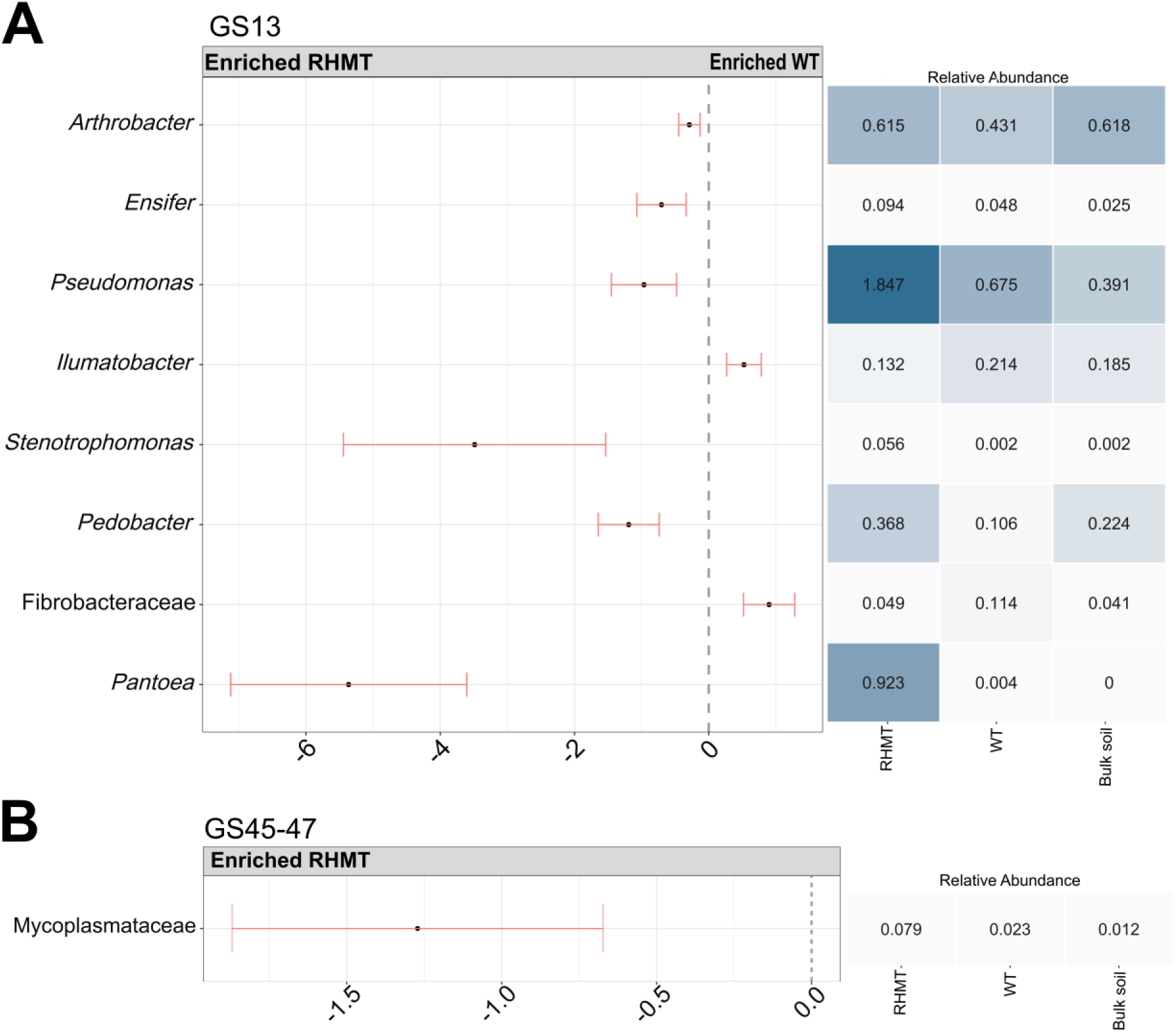
Taxa enriched in the root hair mutant (RHMT) versus the wild-type (WT) across growth stages. At GS11, there were no differentially abundant genera. The differential abundance between the RHMT and WT was determined using beta-binomial regression with corncob. Only taxa that had a p < 0.05 after correcting for multiple testing (FDR) were considered significant. The heatmap displays the mean relative abundances of the taxa in the RHMT, WT, or bulk soil samples.

In summary, the loss of root hair elongation had limited impact on the bacterial community in the wheat rhizosphere, with GS13 being the most affected by the mutation.

### Low-P soil status enriches arbuscule formation and not bacterial P mineralization in response to loss of root hair elongation

To stimulate the root hair mutant to retain P-cycling bacteria further along in its development, we grew the plants to booting (GS45-47) in P-depleted soil. The root hair mutant and wild-type were grown in pots using soil from a historically low-P fertilized plot from a nutrient depletion field trial. In the low-P soil, both genotypes grew only half the biomass compared to when the plants were grown in soil from the P-replete plots of the nutrient depletion field trial (Figure 1 and Figure 4A). The root hair mutant also grew smaller shoots compared to the wild-type in the low-P soil (Figure 4A, Wilcoxon test, p < 0.05). Shoot P concentration of the two genotypes was not different in the low-P soil (Figure 4A, Wilcoxon test, p > 0.05).

**Figure 4.**
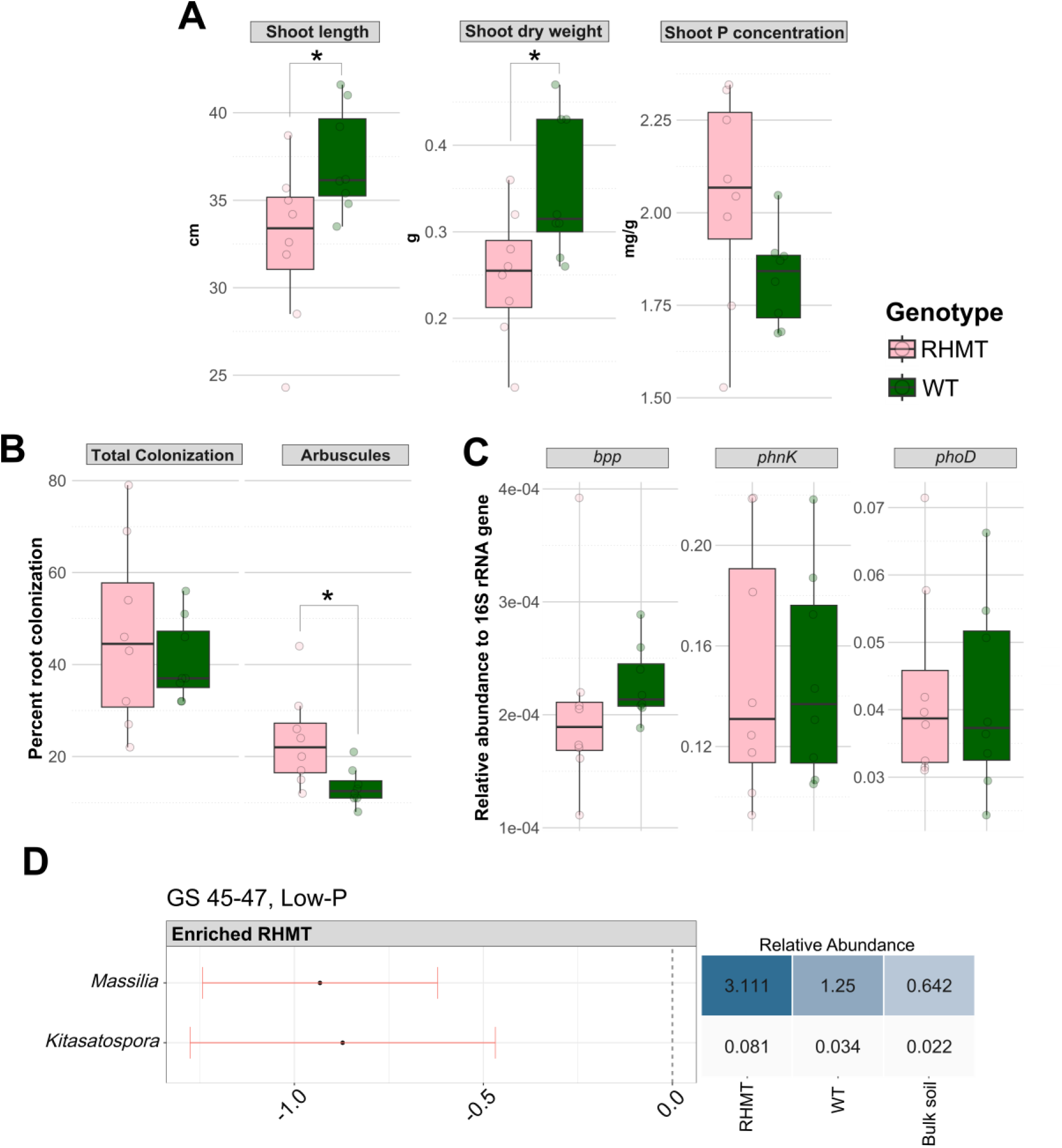
Response of the root hair mutant (RHMT) and wild-type (WT) to low-P conditions at GS45-47. (A) Plant performance as measured by aboveground biomass and shoot P concentration. (B) AMF total colonization and arbuscule formation. (C) Relative abundance of bacterial P mineralization genes. (D) Differentially abundant bacterial genera between the root hair mutant and wild-type. The differential abundance was determined using beta-binomial regression with corncob. Only genera that had a p < 0.05 after correcting for multiple testing (FDR) were considered significant. The heatmap displays the mean relative abundances of the genera in the RHMT, WT, or bulk soil samples. The boxplots display the median, the two hinges (25th and 75th percentile), and the upper and lower whiskers (extending to the largest and smallest values no further than 1.5 times the interquartile range). True data points are overlaid on the boxplots as filled circles. Means were compared with the Wilcoxon test. *p < 0.05.

While the two genotypes were colonized equally by AMF in the low-P soil, the root hair mutant demonstrated higher arbuscule formation (Willcoxon test, p < 0.05) (Figure 4B). The bacterial P mineralization genes that were enriched in the root hair mutant at early seedling growth under P-replete conditions (Table 1) were not enriched under low-P conditions at GS45-47 (Figure 4C). There was an effect of sample source (root hair mutant, wild-type, or bulk soil) on gene relative abundance (one-way ANOVA, p < 0.05, Table S5), but there were no bacterial P-cycling genes enriched in the root hair mutant compared to the wild-type in low-P conditions (Tukey’s HSD, p > 0.05, Table S6).

Soil P status influenced the bacterial microbiome community structure in the rhizosphere (PERMANOVA, p = 0.001, R2 = 0.36), as did the interaction between genotype and soil P status (PERMANOVA, p = 0.014, R2 = 0.07) (Figure S3). Despite this, only two genera, *Massilia* and *Kitasatospora*, were enriched in the root hair mutant under low-P (Figure 4D), and there was no difference in alpha diversity between the two genotypes (Figure S3, Student’s T test, p > 0.05).

In summary, low-P conditions did not increase recruitment of bacterial P-cycling genes in the rhizosphere of the root hair mutant at GS45-47, despite the high P demand at this growth stage. During this growth stage, arbuscule formation was enriched in the root hair mutant under low-P.

## Discussion

The potential of the plant to utilize both the bacterial and fungal fraction of the rhizosphere microbiome for P nutrition has yet to be fully understood. We lack knowledge on the modulation of the P-cycling bacterial community in wheat and how this might be contextualized with regards to AMF, the fungal community vital for plant P nutrition (Smith & Smith, 2011). In this work, we used a recently characterized root hair mutant in wheat (Tsang et al., 2024) to induce a limitation in P uptake (Gahoonia et al., 2001) that can be alleviated by the bacterial microbiome (Zhang et al., 2025). Our results provide novel insights on the importance of bacteria and AMF in compensating for the loss of root hair elongation during wheat development and in P-depleted soils.

With regards to plant development, we found a stunting effect of the *srh1* mutation on early root system development in soil, which was mirrored by diminished aboveground biomass at booting (GS45-47). Reduced root growth is typical of hairless mutants in other crop species grown in soil, highlighting the importance of root hairs in overall crop development (Suzuki et al., 2003; Haling et al., 2013; Ma et al., 2020). Rhizosheath size was also reduced in the root hair mutant, in accordance with previous results on the importance of root hairs on root-soil contact (Haling et al., 2013). Despite potential pleiotropic effects of the *srh1* mutation on crop growth, the inhibition of root hair elongation in the mutant is a useful tool to test our hypotheses on microbiome modulation as a response to the absence of root hair elongation.

In agreement with our first hypothesis, we demonstrated that the bacterial rhizosphere microbiome of the root hair mutant is enriched with P mineralization genes *(bpp, phoD,* and *phnK*) at GS11. Relative abundance of the P solubilization marker gene *pqqC*, the co-factor for gluconic acid production (An & Moe, 2016), was not different between the root hair mutant and wild-type. Since plants rely on root hairs to mobilize organic P into orthophosphate anions (Gahoonia et al., 2001; Richardson et al., 2011; Ma et al., 2020), the specific enrichment of bacterial organic P mineralization genes in the root hair mutant suggests a bacterial route for organic P acquisition to compensate for the loss of root hairs before the AMF symbiosis is established. Indeed, bacteria that metabolize plant root exudates play a significant role in organic P mineralization in the soil (Wang et al., 2025), indicating a potential relationship between microbial function and plant needs. While we cannot rule out competition for P between bacteria and the plant (Raymond et al., 2021), the specific enrichment of mineralization genes may indicate a beneficial relationship over competition.

Our second hypothesis suggested that enrichment of P-cycling genes in the root hair mutant would decrease after AMF symbiosis. Harvest at GS13 revealed that the relative abundance of organic P mineralization genes in the root hair mutant rhizosphere decreased to match that of the wild-type, confirming our hypothesis. This loss of bacterial P mineralization genes at GS13 in the root hair mutant coincided with a marked increase in AMF signal in both genotypes from GS11 to GS13. This aligns with previous work that found AMF colonization in field-grown wheat was absent until GS12 (Marrassini et al., 2024), as the root system required sufficient development prior to initiation of AMF symbiosis (Werner & Kiers, 2015). We also observed that root growth of the mutant, which was stunted in comparison to the wild-type at GS11, increased to match that of the wild-type at GS13. This potentially indicated maturity for AMF colonization or, alternatively, a direct root response to AMF colonization (Zhu et al., 2025). While P uptake in the first weeks of growth plays a vital role in seedling establishment and vigor (Yi et al., 2023), the rate of P uptake in wheat reaches its maximum during booting (Jones et al., 2024). Still, at this late growth stage, the relative abundances of all bacterial P-cycling genes were no different between the mutant and wild-type, while AMF colonization and symbiosis was higher in the root hair mutant. In the absence of root hair elongation, AMF total colonization and arbuscule formation increased, aligning with similar results found in root hair mutants of maize and barely (Jakobsen et al., 2005; Kumar et al., 2019; Ma et al., 2020). Despite the loss of root hairs, which constrains plant P nutrition (Gahoonia et al., 2001), the shoot P concentration was unaffected, indicating dominance of AMF for mineral P uptake. These findings mirrored those found in maize (Kumar et al., 2019).

The time series data across three growth stages in wheat indicated a temporal shift from enrichment of bacterial P-cycling to the dominance of the AMF symbiosis in the absence of root hair elongation. We propose that before the AMF symbiosis is fully dominant, wheat can recruit and rely on bacterial P mineralization to meet their P needs. Once the AMF symbiosis is established, the recruitment of these bacteria is no longer prioritized. Quantifying the plant P stress response across developmental stages, when the crop is grown only with its bacterial rhizosphere microbiome or only in the presence of AMF, would clarify the importance of these microbial fractions in wheat P nutrition. However, since two different soils were used for early and late harvest in this work, direct comparisons of the bacterial P mineralization response to the loss of root hair elongation between these experiments must be carefully interpreted. Ideally, measurements of organic P across the growth stages would provide further insights into how soil organic P conditions may influence the responsiveness of organic P mineralization genes in the rhizosphere of the root hair mutant. The AMF communities, colonization potential, and responsiveness may also be different between the field and the pots (Jansa et al., 2014), and direct parallels between the experiments should be interpreted with restraint.

Alongside functional profiling, long-read 16S rRNA amplicon sequencing revealed the specific taxa responding to the loss of root hair elongation across the three growth stages. However, there was little overlap between the qPCR data and community structure. That is, P-cycling genes only differed between the youngest plants (GS11), while community composition only differed between older plants (GS13 and GS45-47). This disparity was likely due to 16S rRNA sequences that poorly represent functional capabilities (Jaspers & Overmann, 2004), which we have found to be the case for wheat rhizosphere *Pseudomonas* isolates (Herms et al., 2025). Thus, taxonomic profiling is insufficiently sensitive to reveal P-cycling capabilities, and we therefore prioritize the results of the qPCR analyses for interpretation of P-cycling. Nevertheless, we provide evidence for the role of root hairs in shaping the bacterial community in the wheat rhizosphere. We found that the root hair mutant enriched more bacterial taxa compared to its wild-type, as found in barley (Robertson-Albertyn et al., 2017), while maize had an opposite trend (Quattrone et al., 2024). Only one taxon, Actinomycetales, was similarly enriched across wheat and barley root hair mutants (Robertson-Albertyn et al., 2017). Root hair mutants in wheat and maize both enrich the genus *Stenotrophomonas* (Quattrone et al., 2024). *Stenotrophomonas* is a plant-associated bacterium that might play a role in disease suppression, growth promotion, and nutrient acquisition (Kumar et al., 2023). The disruption in microbiome composition in response to the loss of root hair elongation could be attributed to the loss of unique colonization niches (Buddrus-Schiemann et al., 2010; Mousa et al., 2016) or altered root exudate profiles (Quattrone et al., 2024). Analysis of the root exudates of the wheat root hair mutant, while out of scope in this work, could help further clarify the dysbiosis.

Our third hypothesis predicted that we could stimulate the root hair mutant to retain its enriched P-cycling microbiome to booting stage by growing the plants in P-depleted conditions. We speculated that by adding additional P stress, this would initiate a ‘cry for help’ from the plant that would initiate a response from the bacterial microbiome (Zhang et al., 2025). However, this hypothesis was not supported by the data. We did see an enrichment of *Kitatasatospora* in the root hair mutant rhizosphere at GS45-47 in low-P soil, a plant-associated genus that has been indicated in P cycling (Oliveira et al., 2009), but this was not reflected in the functional profiling. The relative abundances of all bacterial P-cycling genes were equal between the root hair mutant and the wild-type. On the other hand, we saw a fungal response in the root hair mutant under these conditions. Total AMF colonization was equal in the root hair mutant in wild-type during booting under low-P conditions, a well-known response of plants grown in P-depleted conditions (Breuillin et al., 2010; Kumar et al., 2019; Zhou et al., 2022). However, arbuscule formation was higher in the root hair mutant compared to the wild-type and increased compared to growth in P-replete conditions. Arbuscules are the specialized mycorrhizal structures that facilitate mineral nutrient translocation from fungus to plant (Smith & Smith, 2011). This result further supports the dominance of AMF in root hair mutants compared to bacteria.

Despite decades of research, successfully manipulating the rhizosphere microbiome for improved crop nutrition remains limited. Our results shed light on how a wheat mutant, defective in root hair elongation, can enrich bacterial and fungal partners for P nutrition. We suggest a temporal component to plant-microbiome interactions that play a role in P nutrition. A practical implication of our findings is that, given genotypic variation in root hair length within elite wheat germplasm (Delhaize et al., 2015), the genotypes with constitutively short root hairs or poor plasticity (Nguyen & Stangoulis, 2019) are likely to enlist microbial mechanisms to ensure P nutrition. Collectively, this work can be used to guide improvements of microbial P uptake mechanisms in wheat. By considering the natural ecology in the rhizosphere of the root hair mutant, we recommend focusing on bacterial interactions only during the early stages of wheat development and otherwise prioritizing the AMF symbiosis throughout wheat growth.

## Acknowledgements

The authors thank Andrew Riche (Rothamsted Research, Harpenden, UK) for access to the high- and low-P managed soils. This study was funded by the Novo Nordisk Foundation (Grant number: NNF19SA0059360) and the Biotechnology and Biological Sciences Research Council (BBSRC) as part of the Collaborative Training Program for Sustainable Agricultural Innovation (CTP-SAI) (grant BB/W009439/1) in partnership with The Morley Agricultural Foundation (TMAF). We thank Stephen Rawsthorne (TMAF) and Jonathan Atkinson (University of Nottingham, UK) for their inputs into IT’s PhD. Further support to CHH was provided by the European Molecular Biology Organization (EMBO) and the Monogram Network.

## Author Contributions

Courtney Horn Herms: design of the research; performance of the research; data analysis, collection, or interpretation; writing the manuscript. Ian Tsang: data analysis, collection, or interpretation. Frederik Bak: data analysis, collection, or interpretation; writing the manuscript. Klara Gunnarsen: data analysis, performance of the research; collection, or interpretation; writing the manuscript. Magdalena Hasenzagl: data analysis, collection, or interpretation. Amaru Miranda Djurhuus: data analysis, performance of the research; collection, or interpretation; writing the manuscript. Tom Thirkell: data analysis, collection, or interpretation; writing the manuscript. Eric Ober: design of the research; data analysis, collection, or interpretation; writing the manuscript. Fiona Leigh: writing the manuscript.

James Cockram: writing the manuscript. Mette Haubjerg Nicolaisen: design of the research; data analysis, collection, or interpretation; writing the manuscript.

## Data Availability

16S rRNA amplicon sequence data is publicly available on NCBI under BioProject PRJNA1308004. Raw data is presented in the figures when appropriate. All other raw data is available upon request.

---

**Supplemental Figure 1.**
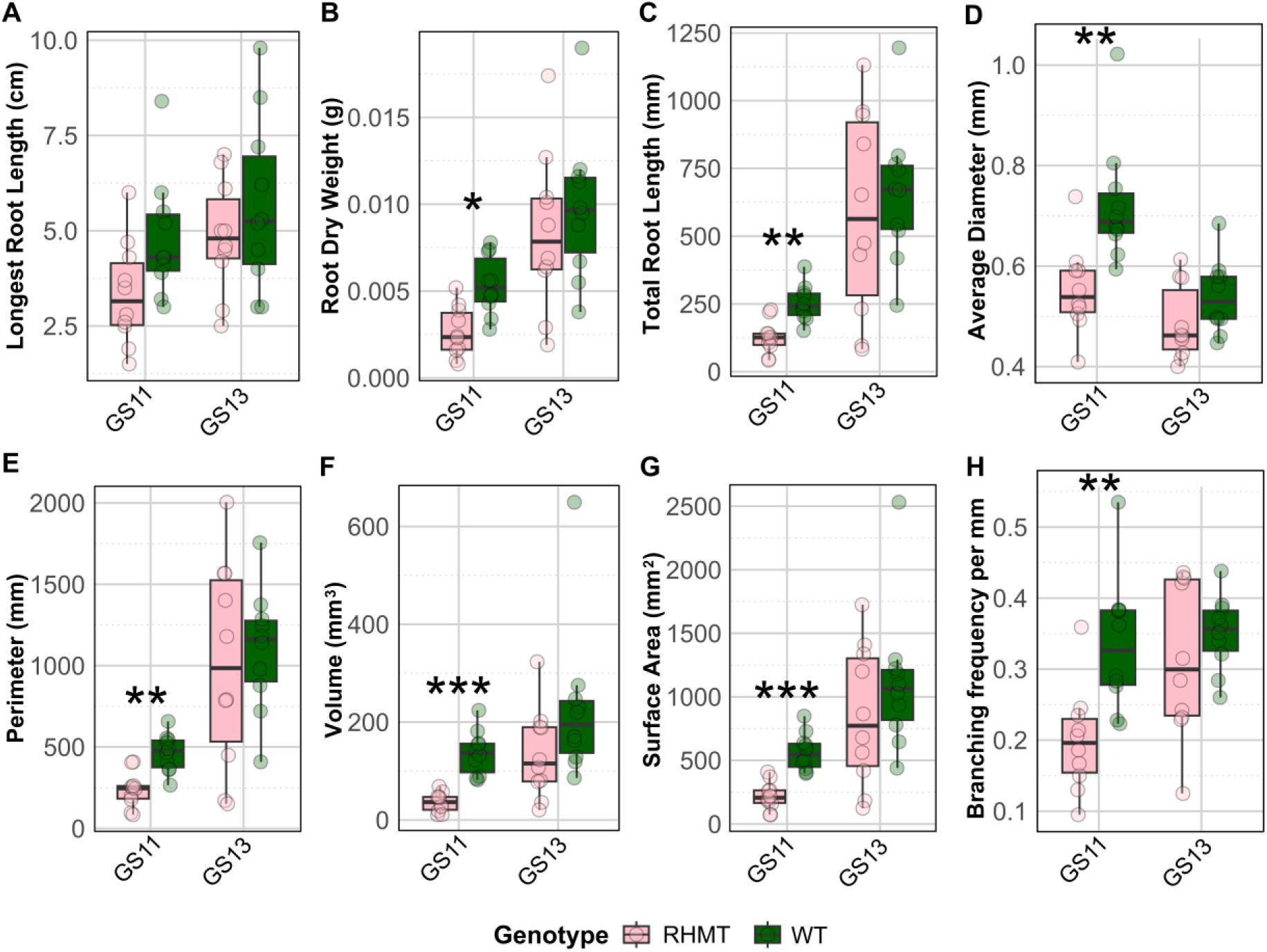
Root phenotypic data of the root hair mutant (RHMT) and wild-type (WT) across the first two growth stages. Genotypes were compared within each growth stage using t-tests (normal data) or Wilcoxon tests (non-normal data). *p < 0.05, **p < 0.01, ***p < 0.001.

**Supplemental Figure S2.**
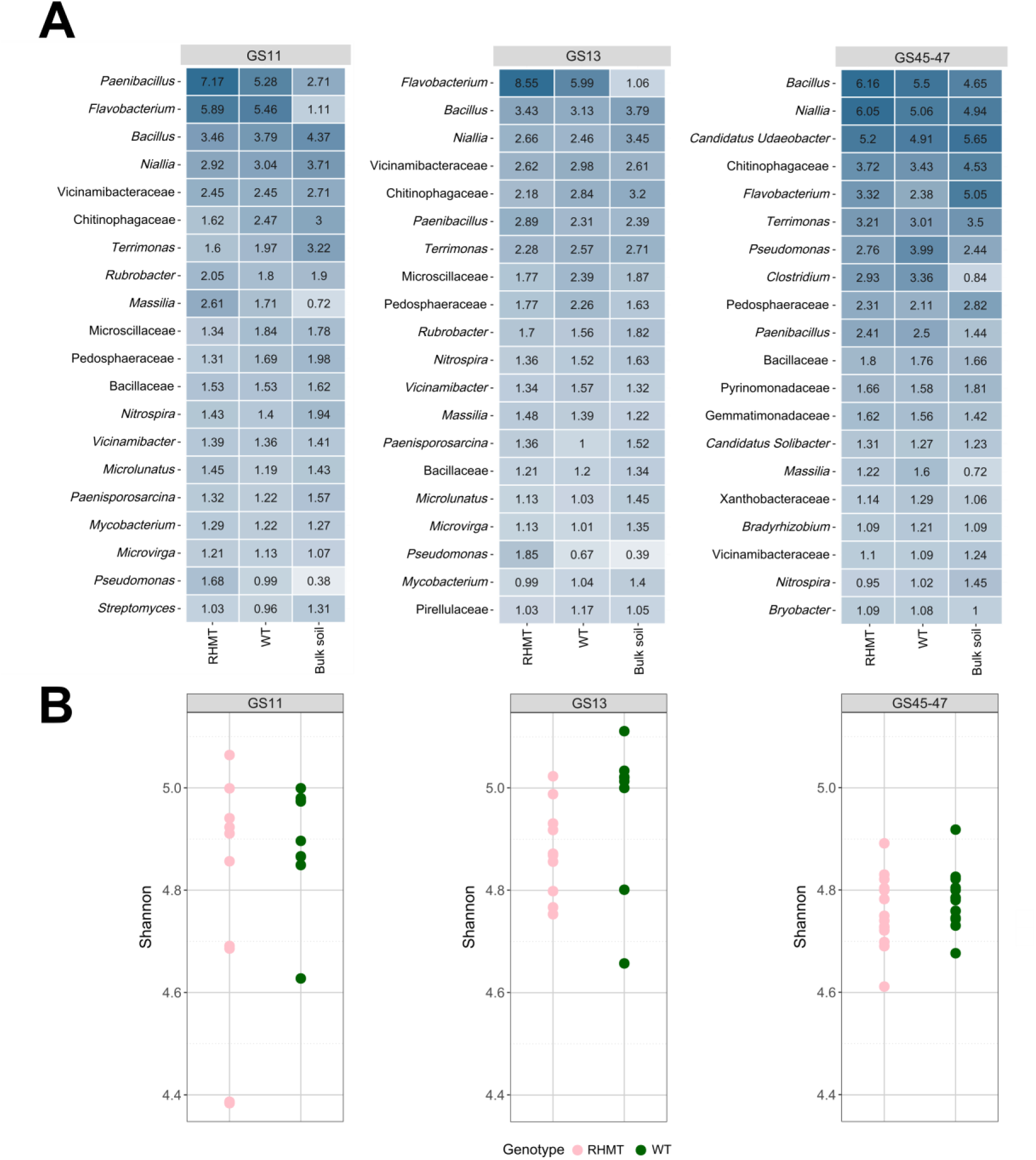
Rhizosphere bacterial community composition of the root hair mutant (RHMT) and wild-type (WT) across growth stages. (A) The heatmap displays the mean relative abundances of the taxa in the RHMT, WT, or bulk soil samples. (B) Alpha diversity as described by the Shannon diversity index across growth stages in the RHMT and WT. Each point represents a sample. Genotypes were compared within each growth stage using t-tests (normal data) or Wilcoxon tests (non-normal data). *p < 0.05.

**Supplemental Figure S3.**
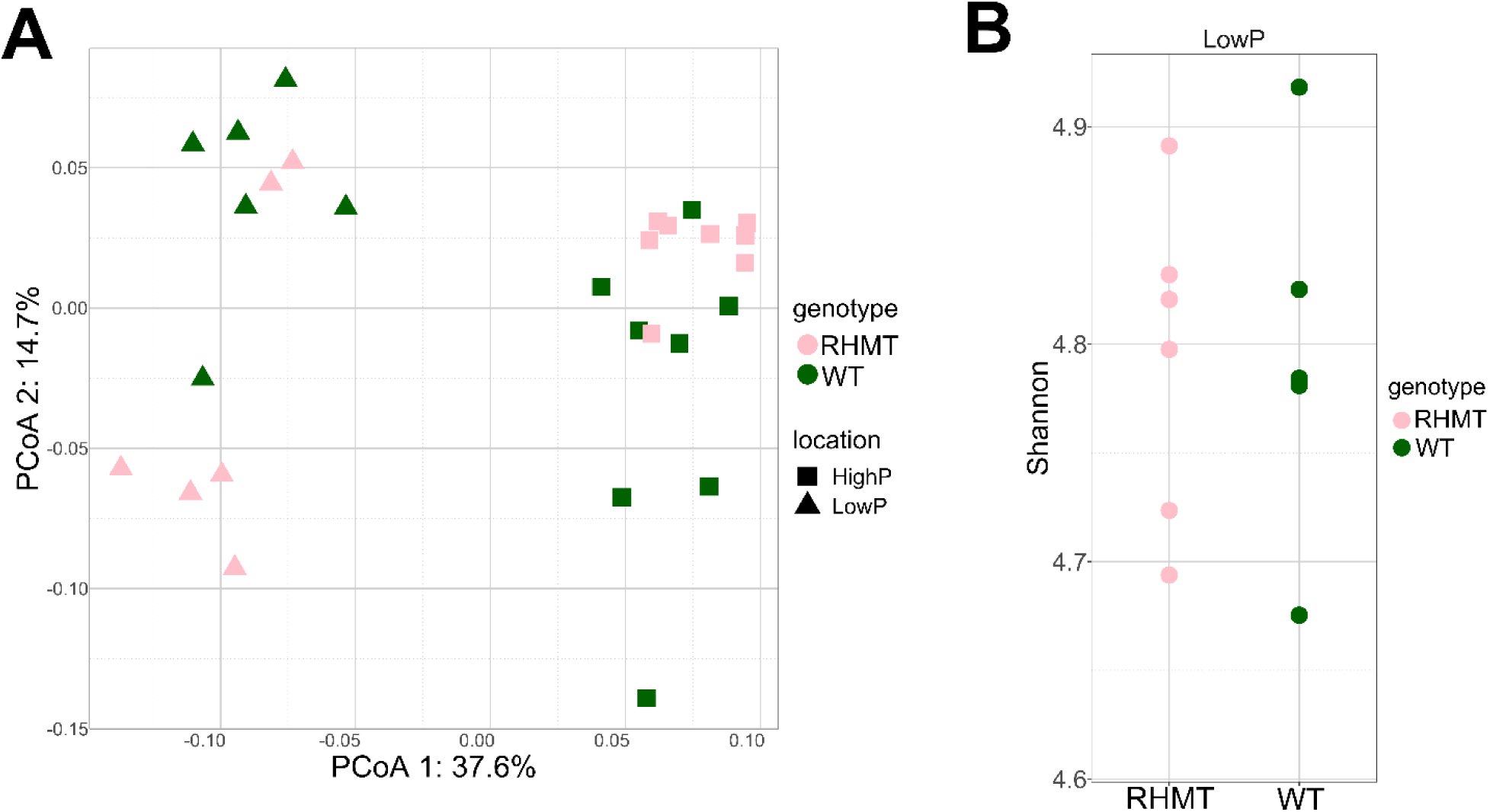
Bacterial community composition of the rhizosphere of the root hair mutant (RHMT) and wild-type (WT) under high- and low-P soil conditions. (A) Principal coordinate analysis (PCoA) plot based on Bray-Curtis dissimilarities. Each point represents a sample. (B) Alpha diversity as described by the Shannon diversity index. Each point represents a sample.

